# New insights into the plasma and urinary metabolomic signatures of spontaneously hypertensive rats

**DOI:** 10.1101/2025.02.28.640925

**Authors:** Celia Rodríguez-Pérez, Alejandra Vázquez-Aguilar, Oscar D. Rangel-Huerta, Estefanía Sánchez-Rodríguez, Ángel Gil, Caridad Díaz, Félix Vargas, María D. Mesa

**Affiliations:** Department of Nutrition and Food Science, Faculty of Pharmacy, University of Granada, Campus of Cartuja, 18071 Granada, Spain; Institute of Nutrition and Food Technology (INYTA) ‘José Mataix’, Biomedical Research Centre, University of Granada, Avda. del Conocimiento s/n, 18071 Granada, Spain; Instituto de Investigación Biosanitaria ibs. GRANADA, 18012 Granada, Spain; Department of Biochemistry and Molecular Biology II, University of Granada, Campus of Cartuja, 18071 Granada, Spain; Section of Chemistry and Toxinology, Norwegian Veterinary Institute. P.O. Box 64, N-1431 Ås, Norway; CIBEROBN (CIBER Physiopathology of Obesity and Nutrition CB12/03/30028), Institute of Health Carlos III (ISCIII), Madrid, Spain; Department of Screening & Target Validation, Fundación MEDINA, 18016 Granada, Spain; Department of Physiology, Faculty of Medicine, University of Granada, Parque Tecnológico de la Salud, Avenida de la Investigación s/n, 18100 Armilla, Granada, Spain

**Keywords:** Hypertension, SHR rats, metabolomics, urine, plasma, *mummichog* pathway enrichment

## Abstract

**BACKGROUND:** Hypertension is a major risk factor associated with cardiovascular diseases and one of the leading causes of premature death. Hypertension has no obvious clinical signs; therefore, the identification of new biomarkers for the early detection of people at risk is needed to prevent hypertension and its further complications. Metabolomics is a useful tool for studying in vivo metabolic profiles that better understand the pathogenesis of diseases such as hypertension. This work aimed to explore the plasma and urinary non-targeted metabolic profile of 16-week-old spontaneously hypertensive rats (SHR) to identify new metabolomic profiles associated with hypertensive phenotypical characteristics that allow early diagnosis of patients at risk.

**METHODS:** Plasma and 24-hours urine samples were collected from 10 SHR and 10 age-matched normotensive Wistar-Kyoto 16-week-old male rats. Plasma and urinary metabolic profiles were investigated using high-performance liquid chromatography quadrupole time of flight coupled to mass spectroscopy followed by multivariate statistical analysis. The *mummichog* pathway enrichment analysis was used to integrate metabolomics data into biological contexts.

**RESULTS:** A total of 16 differential metabolites were found in plasma and 13 differential metabolites in urine from SHR compared to normotensive rats. Differences in some microbiota-derived metabolites suggest changes in the gut microbiota associated with hypertension in our experimental model. The *mummichog* algorithm has recognized that hypertensive metabolism is associated with the altered metabolism of steroid hormones, bile acid and purines.

**CONCLUSIONS:** This work highlights the importance of metabolomics as a tool for the identification of biomarkers in the early detection of hypertension, a condition that represents a significant risk to cardiovascular health. The findings suggest that hypertension is associated with alterations in the metabolism of steroid hormones, bile acids and purines, as well as metabolites derived from the intestinal microbiota. These results not only contribute to a better understanding of the pathogenesis of hypertension but also open the door to future research in humans that could facilitate more accurate and earlier diagnoses, thus helping to prevent serious complications associated with this disease.

## 1. INTRODUCTION

Arterial hypertension is a complex multifactorial vascular pathology caused by the interactions between environmental and polygenic factors. This disease is associated with other adverse cardiovascular complications, including myocardial infarction, stroke, kidney disease, and global mortality. It has been reported as one of the leading causes of premature death^1^, and it is the world’s most prevalent cardiovascular disorder, affecting 1.28 billion adults aged 30–79 years worldwide. However, as hypertension has no obvious clinical signs, it has been estimated that approximately half of the population with hypertension (41% of women and 51% of men) is not aware of their condition; therefore, they are not treated^2^. Hence, hypertension is considered a silent killer^3^. In addition, despite the availability of several preventive and therapeutic approaches, arterial hypertension continues to be an unresolved risk factor for disease burden worldwide^4^. Therefore, understanding the mechanisms related to the pathogenesis of hypertension and the onset of its complications is essential for identifying new biomarkers for early detection to prevent hypertension and its complications.

In this regard, *in vivo* models of hypertension are crucial to completely understand the mechanisms involved and to develop novel therapeutic approaches that can be applied to human research. Among different preclinical hypertensive animal models, spontaneously hypertensive rats (SHR) are the most widely used genetic models of hypertension for the study of primary or essential hypertension due to their similarities to human hypertension^5,6^. The systolic blood pressure (SBP) of SHR reaches 180–200 mm Hg after 4 weeks of growth, while their breeding brother Wistar-Kyoto rats (WKY) remain normotensive. The SHR model has been used to identify hypertension-related genes, to evaluate targeted organ complications, and for the screening of potential pharmacological drugs; indeed, they are expected to provide insight into pathological and therapeutic mechanisms for the regulation of blood pressure. Interestingly, hypertension has also been linked to immune disorders and dysbiosis^7^, but the molecular mechanisms have not been fully elucidated and require further investigation^8^.

Metabolomics is an emerging discipline that characterizes the small molecules (< 2000 Da) present in a biological sample that are closely related to an organism’s phenotype, providing wide information about biological systems and metabolism. However, it requires the use of robust bioinformatics techniques and statistical strategies to mine large amounts of data to extract relevant biological information, as well as other novel tools, such as *mummichog,* oriented to predict metabolic patterns that may be associated with a disease, rather than individual identification of metabolites^9^.

Although earlier studies have reported a number of metabolites associated with the development of hypertension in SHR^10^ and humans^8^, which depend on the stage of the disease, the exact implications of the metabolic pathways involved in cardiac dysfunction have not yet been well established^11^. Therefore, the present study aimed to compare *in vivo* plasma and urine metabolic differences between hypertensive SHR and normotensive healthy WKY rats, using liquid chromatography-mass spectrometry non-targeted metabolomic strategies, to identify possible metabolic pathways implicated in hypertension that may be selected as possible therapeutic targets.

## 2. SUBJECTS AND METHODS

### 2.1. Animals and study design

A total of 20 (n=10 SHR and n=10 Wistar-Kyoto (WKY)) 16-week-old male rats were employed in this study. The animals were purchased from Janvier Labs (CEDEX, France). All the rats had *ad libitum* access to food and water. For adaptation to housing conditions, the animals were fed a standard maintenance diet (Panlab) containing barley, wheat, maize, soybean meal, wheat bran, hydrolyzed fish proteins, dicalcium phosphate, a pre-mixture of minerals, calcium carbonate, a pre-mixture of vitamins, and 68.9% carbohydrates (fiber 3.9%), 16.1% proteins, and 3.1% fat. The experiment was performed in accordance with the guidelines set by the European Community Council Directives for the Ethical Care of Animals (86/609/EEC) and was approved by the Ethics Committee of Laboratory Animals of the University of Granada (Spain, permit number 18/07/2017/099).

SBP was monitored during the adaptation period to evaluate disease evolution. SBP was measured by plethysmography in conscious rats (LE 5001-Pressure Meter, Letica SA, Barcelona, Spain); at least seven determinations were made at every session, and the mean of the lowest three values within a range of 5 mmHg was set as the final SBP value. Food and drink intake and body weight were also monitored. At the end of the maintenance period, the rats were introduced into metabolic cages (Panlab, Barcelon, Spain) for 24-h urine collection. Fasting rats were anesthetized with 2.5 ml per kg equitensin (i.p.) and blood was drawn via abdominal aortic puncture. Blood was centrifuged at 1750 × g for 10 min at 4°C. Aliquots of plasma were frozen immediately at -80°C until metabolomic analysis.

### 2.2. Metabolite extraction

Plasma and urine samples were thawed on ice and kept at 4°C during the entire extraction process and pretreated following the method proposed by Liu et al. with some modifications^11^. Briefly, 100 μl of plasma samples were deproteinized with 200 μl of a precipitation mixture containing acetonitrile, methanol, and acetone (8:1:1 v/v). The samples were vortexed and placed at -20°C for 30 min to facilitate protein precipitation. The samples were centrifuged at 14800 rpm for 10 min at 4°C, and the supernatants were evaporated in an Eppendorf^®^ Concentrator Plus evaporator centrifuge for 2 h. Dried samples were reconstituted in 100 μl of 0.1% formic acid in water, vortexed for 20 s, and incubated in an ice bath for 10 min. Subsequently, the samples were centrifuged at 14800 rpm for 10 min at 4°C. Finally, 40 μl of the supernatant was transferred to a high-performance liquid Chromatography (HPLC) vial with 250 μl inserts, to which 5 μl of internal standard (IS), consisting of a mixture of roxithromycin, tryptophan, hydroxydiclofenac, warfarin, ibuprofen, and bisphenol at different concentrations, was added. The IS allowed the monitoring of instrument performance and aided chromatographic alignment. The prepared samples were stored at -80 °C until their analysis.

In the case of urine, after measuring the osmolarity (Gonotec Osmomat 030 D Osmometer) and adjusting for differences in hydration status among animals, all samples were diluted to 200 ± 20 mOsm/kg using Milli-Q water^12^. The samples were then centrifuged at 14800 rpm for 10 min at 4°C, 40 μl of the supernatant was transferred to an HPLC vial, and 5 µL of the abovementioned IS was added. The prepared samples were stored at -80 °C until their analysis.

Quality control (QC) samples from plasma and urine were prepared by pooling equal volumes (20 μl) of all studied samples. Blanks were prepared using water.

### 2.3. HPLC-ESI-QTOF-MS Analysis

Samples were analyzed using an Agilent Series 1290 High Performance Chromatography (Agilent Technologies, Santa Clara, CA, USA) coupled to an AB SCIEX TripleTOF 5600 quadrupole time-of-flight mass spectrometer (qTOF-MS). Chromatographic separation was performed using a Waters Atlantis T3 C18 chromatographic column (2.1 mm x 150 mm, 3 µm (Waters Corporation, Milford, MA, USA), which was maintained at 30°C in the oven. The mobile phases were 0.1% formic acid in a mixture of water (90:10) (A) and 0.1% formic acid in MeCN: water (90:10) (B). The column was eluted with the following gradient: 0 to 0.5 min, 0% eluent B, 0.5 to 11 min, 100% eluent B, which was maintained until 15.60 min and 0% of eluent B and 15.60-20 min 0% eluent B at a constant flow rate of 0.3 mL/min. The injection volume was 5 µl.

Triple TOF 5600 operating in positive- and negative-ion modes was employed for metabolite detection using a mass range of 50–1250 Da. The Triple TOF used a Duo Spray source with separate electrospray ionization (ESI) and atmospheric-pressure chemical ionization probes. ESI was used to analyze the metabolomic profile of the sample. The ESI parameters were as follows: positive mode, capillary voltage of 5000 V, nebulizer gas pressure of 50 psi, drying gas pressure of 50 psi, temperature of 500°C, and focusing potential of 100 V. Negative mode: capillary voltage -4500 V, nebulizer gas pressure, 50 psi; drying gas pressure, 50 psi; temperature, 500 °C; focusing potential, 100 V. Both analyses were performed using independent data acquisition, and the eight most intense ions in every cycle were fragmented.

To avoid possible bias due to sequential injection within the same group in the LC-qTOF-MS chromatographic profile, the sample injection sequence was randomized. QC samples were injected at the start of the analysis and every five studied samples aiming to stabilize the instrument conditions and monitor the system performance. Additionally, QC samples were used to adjust the signal intensity drift within and between batches.

### 2.4. Data acquisition and analysis

The MarkerView software (version 1.2.1, AB SCIEX, Concord, ON) was employed for data set creation with the following extraction parameters: retention time (RT) 1.20-15.00 min; offset subtraction 10 scans; Subtraction Mult. Factor 1.3, noise floor 50 counts per second (cps), minimum spectral peak width 0.02 Da; minimum peak width RT (4 scans); RT tolerance (0.12 min); mass tolerance (10 ppm); required number of samples (6), and maximum number of peaks 5000. After aligning the data, anomalous results were filtered, and a data matrix was obtained containing information on mass data, RT, and peak areas of normalized sample spaces, blanks, and QC for each of the plasma and urine samples tested in positive and negative ionization modes.

### 2.5. Data preprocessing

From the raw data matrix obtained in the previous step, another matrix was extracted with the RT, the mass-charge ratio (m/z), and the areas of metabolic characteristics detected in the filtered data. Alternative non-parametric measures of the relative standard deviation and the D-ratio were calculated with a cut-off point of 0.2 and 0.4, respectively. In the case of missing values, imputation was performed using the Random Forest algorithm. All calculations were performed using the Notame package in R (v0.0.900)^13^. Subsequently, the data was evaluated to identify technical drift, which was corrected by modeling using a spline-cubic regression based on the QC samples and then corrected for the abundance of all samples by reversing the modeling effect of the drifts. This process was developed independently for each feature.

### 2.6. Multivariate and univariate statistical analyses

Multivariate statistical analyses were performed using the SIMCA-P software (V16; Sartorius Stedim Biotech, Umeå, Sweden) following the workflow described by Rangel-Huerta et al.^14^. Datasets were scaled using Pareto scaling. Principal Component Analysis (PCA) was employed to assess the quality of metabolomic data on a QC basis to discard potential outliers, identify clustering patterns, and visualize the total variation of the metabolite profiles. Orthogonal Partial Least Squares Discriminant Analysis (OPLS-DA) was used to identify metabolite patterns and specific features that discriminate between groups. A seven-round cross-validation was applied to this model, and subsequently, the cross-validation analysis of variance (CV-ANOVA) was calculated to assess the reliability of the models. A p-value ≤ 0.05 was considered significant. Additionally, the R2X and Q2 were evaluated to assess robustness, considering that values close to one reflect a reliable model. Variable selection of the most relevant features was performed using an S-plot, considering a p > 0.05, p(corr) > 0.5, and an adjusted p-value from a t-test (p < 0.05) was considered significant.

### 2.7. Pathway analysis and metabolite identification

The MS Peaks to Pathways module from MetaboAnalyst was employed to predict pathway activity from untargeted mass spectral data was employed^15^. To this end, *p*-values and *t*-scores were determined for all metabolites from both the plasma and urine samples of the SHR and WKY groups. The analysis was performed in mixed mode (positive and negative) with the following parameters: 1) instrument mass precision of 10 ppm, 2) retention time (RT) in minutes, 3) p-value, 4) positive/negative analytical mode, and 5) *Mummichog* algorithm with a cut-off value of p ≤ 0.05 and also the gene set enrichment analysis (GSEA) activated. To minimize false positives, pathways with an overlap size smaller than three were excluded from the analysis. *R. norvegicus* was selected from the KEGG pathway map for the analysis. Pathways with a combined p-value (*Mummichog* and GSEA) ≤ 0.1 were considered significant.

Subsequently, KEGG GENOME (<https://www.genome.jp/kegg/kegg2.html>) was accessed using the codes of the compounds previously identified in MetaboAnalyst by applying a filter of sets of metabolites associated with the route, and the chemical compounds were annotated in this analysis.

The molecular annotation of the relevant compounds obtained in the multivariate analysis was carried out in SIRIUS 4.9.12^16^ and public databases using MS-DIAL software^17^. The identity of the metabolites of interest was confirmed by considering the mass accuracy and true isotopic pattern in both the MS and MS/MS spectra provided by qTOF-MS.

## 3. RESULTS

### 3.1. Animal description

The SBP, heart weight, and left ventricular weight were higher in SHR *vs* WKY rats, as well as water intake and diuresis, and heart rate tended to be higher (p=0.054) (Table 1). The 24-h food intake was similar in SHR, as well as tibia length and kidney weight, while the body and liver weights were lower in SHR than in WKY rats (Table 1).

**Table 1.**
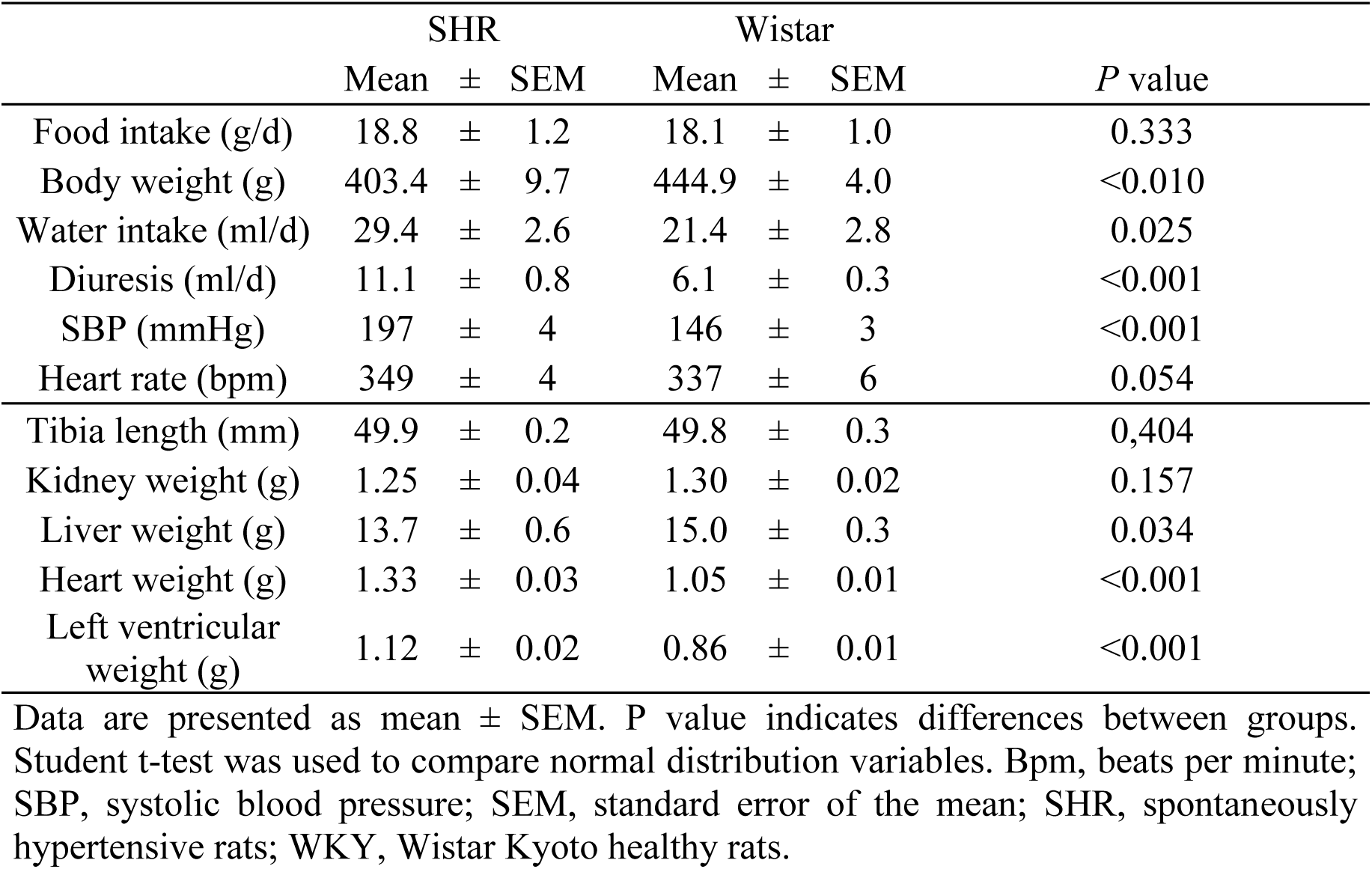
Food and water intakes, body weight, diuresis, SBP and heart rate, and the tibia length, kidney weight, liver weight, heart weight, and the left ventricular weight, of the 16 weeks-old SHR and control Wistar-Kyoto rats.

|  | SHR |  |  | Wistar |  |  | *P* value |
| --- | --- | --- | --- | --- | --- | --- | --- |
|  | Mean | ± | SEM | Mean | ± | SEM |  |
| Food intake (g/d) | 18.8 | ± | 1.2 | 18.1 | ± | 1.0 | 0.333 |
| Body weight (g) | 403.4 | ± | 9.7 | 444.9 | ± | 4.0 | <0.010 |
| Water intake (ml/d) | 29.4 | ± | 2.6 | 21.4 | ± | 2.8 | 0.025 |
| Diuresis (ml/d) | 11.1 | ± | 0.8 | 6.1 | ± | 0.3 | <0.001 |
| SBP (mmHg) | 197 | ± | 4 | 146 | ± | 3 | <0.001 |
| Heart rate (bpm) | 349 | ± | 4 | 337 | ± | 6 | 0.054 |
| Tibia length (mm) | 49.9 | ± | 0.2 | 49.8 | ± | 0.3 | 0.404 |
| Kidney weight (g) | 1.25 | ± | 0.04 | 1.30 | ± | 0.02 | 0.157 |
| Liver weight (g) | 13.7 | ± | 0.6 | 15.0 | ± | 0.3 | 0.034 |
| Heart weight (g) | 1.33 | ± | 0.03 | 1.05 | ± | 0.01 | <0.001 |
| Left ventricular weight (g) | 1.12 | ± | 0.02 | 0.86 | ± | 0.01 | <0.001 |

Data are presented as mean ± SEM. *P* value indicates differences between groups. Student t-test was used to compare normal distribution variables. Bpm, beats per minute; SBP, systolic blood pressure; SEM, standard error of the mean; SHR, spontaneously hypertensive rats; WKY, Wistar Kyoto healthy rats.

### 3.2. LC-qTOF-MS untargeted metabolomics analysis

After pre-processing, quality assessment, data filtering, and clustering, a dataset containing 195 features in the positive mode and 174 in negative mode for plasma and 162 features in the positive mode and 239 in the negative mode for urine was used for further statistical analysis.

An unsupervised multivariate statistical approach was used to compare the metabolomic differences between the SHR and WKY rats. The PCA score plots of plasma and urine (Figure 1A and 1B, respectively) indicated good separation between the SHR and WKY animal models. The score plot for the first two principal components does not reveal any outliers.

**Figure 1.**
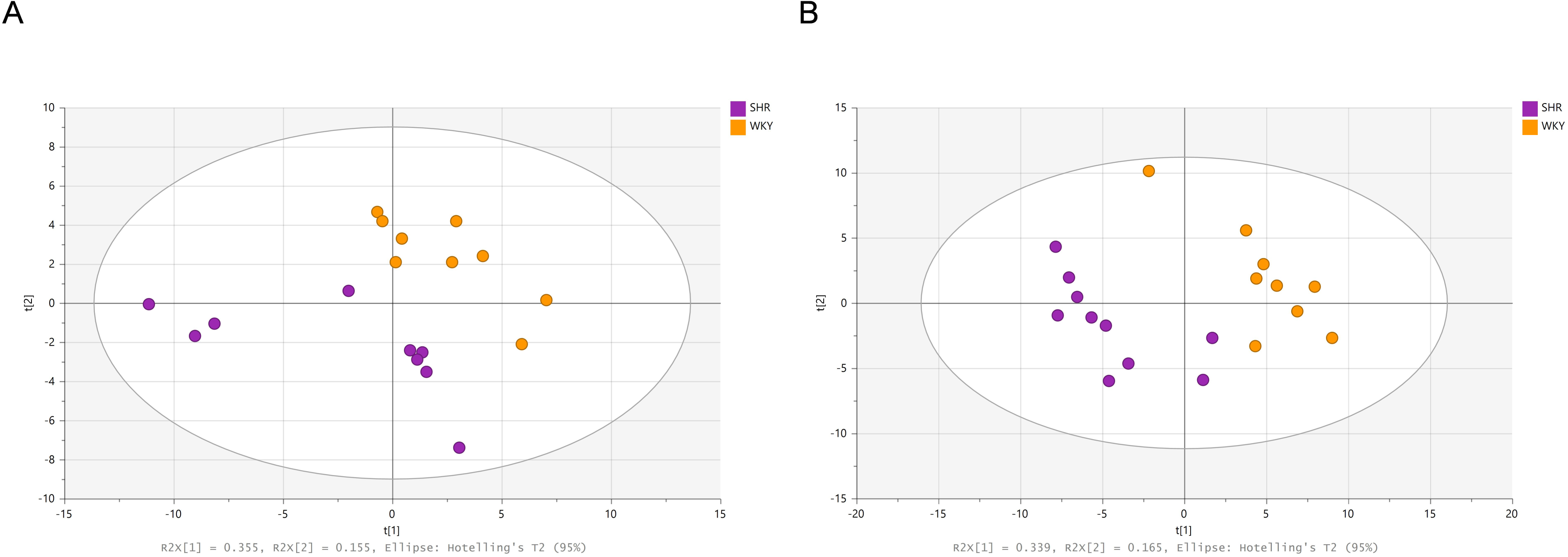
PCA scores plot from LC-qTOF-MS data for the plasma (A) and urine (B) of SHR and WKY animal models. Separation along the Y-axis represents variation between animal models. LC-qTOF-MS, Liquid Chromatography coupled to quadrupole time-of-flight mass spectrometer. PCA, principal component analysis; SHR, spontaneously hypertensive rats; WKY, Wistar Kyoto healthy rats.

The supervised multivariate analysis OPLS-DA indicated a strong separation between the two groups (data not shown). One hundred permutation tests were conducted to evaluate the statistical robustness of the plasma and urine OPLS-DA models. Good discrimination was observed between the two groups with values obtained for the plasma model: R^2^X (0.504), R^2^Ycum (0.954), Q^2^cum (0.874), and CV-ANOVA of 9.38E-06, and for the urine model: R^2^X (0.572), R^2^Ycum (0.977), Q^2^cum (0.837), and CV-ANOVA of 0.00253, thus indicating that the models were reliable and provided excellent prediction ability. Additionally, the S-plot derived from the OPLS-DA models depicts a scatter plot that combines the modeled covariance (X-axis) and modeled correlation (Y-axis) from the OPLS-DA, allowing the identification of interesting variables. Figures 2A and 2B show the plasma and urine S-plots, respectively. The variables showing p[1] > 0.05 and p(corr) > 0.5 values were considered the most relevant metabolites for the differentiation between samples.

**Figure 2.**
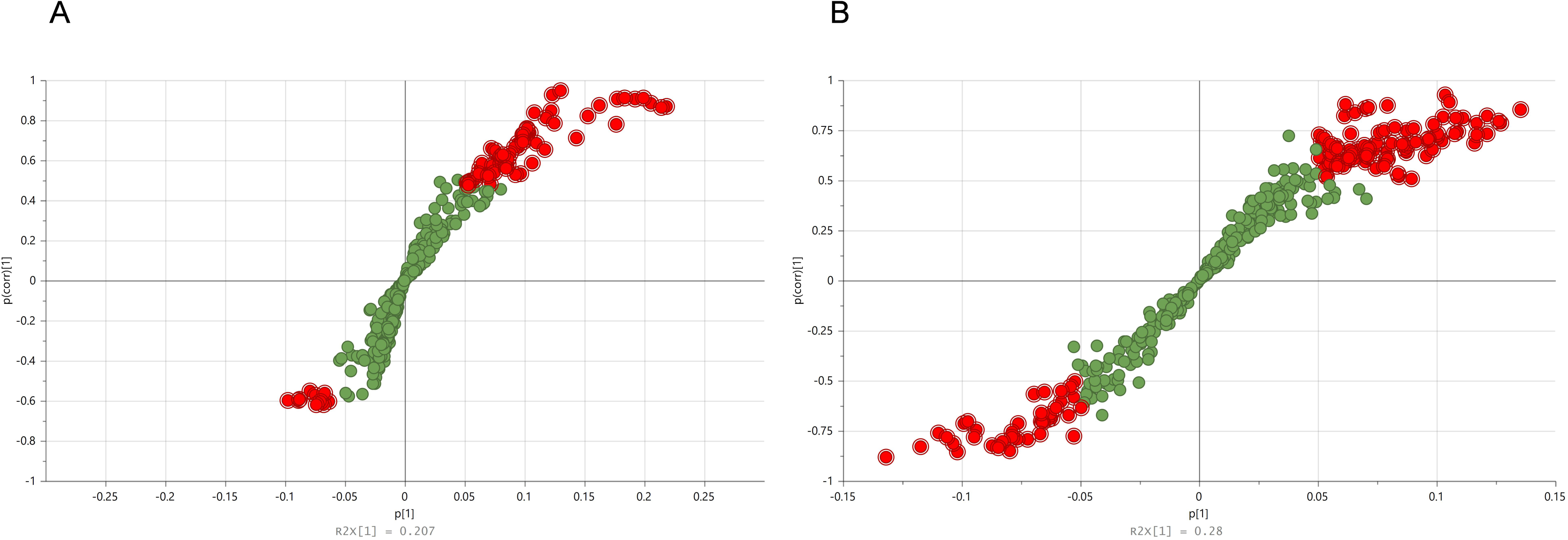
S-plot derived from the OPLS-DA model indicating plasma (A) and urine (B) biomarkers increased in WKY (upper right) and SHR (lower left). Variables with a p > 0.05 and p(corr) > 0.5 were considered significant and are marked in red. OPLS-DA, orthogonal partial least squares discriminant analysis; SHR, spontaneously hypertensive rats; WKY, Wistar Kyoto healthy rats.

Specific observations were made on the differential metabolites identified in the SHR compared with those in the WKY rats. Of the 59 and 111 differential metabolites in plasma and urine, respectively, 16 were annotated in plasma and 14 in urine. These metabolites are presented in Tables 2 and 3, respectively, indicating their RT, observed m/z, suggested ion, fragments from MS/MS experiments, probability, p(corr) value from S-plots, annotation, and family class.

**Table 2.** Plasma identified discriminant metabolites between SHR and WKY rats.

| RT<br>(min) | Observed<br>*m/z* | Suggested ion | Fragments | Probability* | p(corr) | Suggested annotation | Family class |
| --- | --- | --- | --- | --- | --- | --- | --- |
| **Negative ionization mode** |  |  |  |  |  |  |  |
| 4.83 | 283.083 | [M – H]<sup>–</sup> | 75.0077, 113.0226, 85.0293,<br>57.0337 | 100% | 0.67 | p-Cresol glucuronide | Organic acids and derivatives |
| 6.54 | 187.007 | [M – H]<sup>–</sup> | 107.0500 | 100% | 0.06 | p-Cresol sulfate | Organic acids and derivatives |
| 8.75 | 407.280 | [M – H]<sup>–</sup> | 389.2711, 345.2808 | 100% | 0.80 | Cholic acid | Steroids and derivatives |
| 10.26 | 538.313 | [M + CH<sub>2</sub>O<sub>2</sub> - H]<sup>–</sup> | 478.2946, 253.2185 | 98% | 0.75 | LPC (16:1) | Glycerophospholipids |
| 10.69 | 524.278 | [M – H]<sup>–</sup> | 327.2322, 283.2428, 214.0489 | 93% | 0.61 | LPE(22:6) | Glycerophospholipids |
| 11.04 | 552.331 | [M + CH<sub>2</sub>O<sub>2</sub> - H]<sup>–</sup> | 492.3098, 267.2335 | 99% | 0.70 | 1-(10Z-heptadecenoyl)-sn-glycero-3-phosphocholineLPC(17:1) | Glycerophospholipids |
| 11.32 | 590.349 | [M + CH<sub>2</sub>O<sub>2</sub> - H]<sup>–</sup> | 530.3276, 305.2507 | 90% | 0.60 | 1-(5Z,8Z,11Z-eicosatrienoyl)-sn-glycero-3-phosphocholine | Glycerophospholipids |
| **Positive ionization mode** |  |  |  |  |  |  |  |
| 8.77 | 426.321 | [M + H<sub>3</sub>N + H]<sup>+</sup> | 355.2643, 373.2751 | 100% | 0.82 | Cholic acid | Steroids and derivatives |
| 9.83 | 468.309 | [M + H]<sup>+</sup> | 184.0743, 450.2996, 104.1072 | 100% | 0.64 | LPC (14:0) | Glycerophospholipids |
| 10.04 | 494.324 | [M + H]<sup>+</sup> | 184.0743 | 91% | 0.78 | LPE (16:1) | Glycerophospholipids |
| 10.06 | 542.326 | [M + H]<sup>+</sup> | 184.0736, 524.3126, 104.1066 | 96% | 0.60 | LPC (20:5) | Glycerophospholipids |
| 10.26 | 494.323 | [M + H]<sup>+</sup> | 476.3156, 184.0746, 104.1080 | 99% | 0.74 | LPC(16:1) | Glycerophospholipids |
| 11.04 | 508.342 | [M + H]<sup>+</sup> | 184.0729, 490.3279 | 98% | 0.72 | LPC(17:1) | Glycerophospholipids |
| 11.53 | 546.354 | [M + H]<sup>+</sup> | 184.0746, 528.3466, 104.1076 | 94% | 0.63 | LPC (20:3) | Glycerophospholipids |
| 11.72 | 480.309 | [M + H]<sup>+</sup> | 339.2902, 462.2994, | 100% | 0.67 | LPE (18:1) | Glycerophospholipids |
| 12.03 | 572.372 | [M + H]<sup>+</sup> | 184.0738, 554.3624, 104.1072 | 92% | 0.65 | LPC (22:4) | Glycerophospholipids |

LPC, lysophosphatidylcholines; LPE, Lysophosphatidylethanolamine; *m/z* mass to charge ratio; PE, phosphatidylethanolamine; RT retention time. \*Corresponds to class assignment. Only those compounds that had MS/MS information have been included.

**Table 3.** Urine identified discriminant metabolites between SHR and WKY rats.

| RT (min) | Observed $m/z$ | Suggested ion | Fragments | Probability* | p(corr) | Suggested annotation | Family class |
| --- | --- | --- | --- | --- | --- | --- | --- |
| **Negative ionization mode** |  |  |  |  |  |  |  |
| 1.43 | 243.062 | [M – H]<sup>-</sup> | 183.0410, 124.0259 | 98% | -0.66 | Pseudouridine | Nucleoside and nucleotide analogues |
| 1.47 | 167.0211 | [M – H]<sup>-</sup> | 124.0148 | 99% | -0.79 | Uric acid | Organoheterocyclic compounds |
| 1.65 | 191.0198 | [M – H]<sup>-</sup> | 173.0090, 103.0397 | 94% | -0.52 | Citric acid | Organic acids and derivatives |
| 2.58 | 357.0839 | [M – H]<sup>-</sup> | 339.0720, 181.0502,<br>175.0250, 137.0602,<br>113.0237 | 100% | 0.65 | DHPPA (A-D-glucuronide) | Organic oxygen compounds |
| 3.55 | 326.0894 | [M – H]<sup>-</sup> | 175.0250, 150.0560,<br>108.0447, 113.0235 | 100% | 0.50 | Unknown | Organic oxygen compounds |
| 4.36 | 178.0518 | [M – H]<sup>-</sup> | 160.0394, 148.0406,<br>134.0613, 77.0393 | 100% | 0.76 | Hippuric acid | Benzenoids |
| 4.78 | 193.0513 | [M – H]<sup>-</sup> | 178.0280, 149.0610,<br>134.0376 | 96% | 0.76 | Isoferulic acid | Phenylpropanoids and polyketides |
| 4.78 | 273.0091 | [M – H]<sup>-</sup> | 193.0507, 178.0275,<br>149.0608, 134.0371 | 97% | 0.80 | Ferulic acid-4-O-Sulfate | Phenylpropanoids and polyketides |
| 4.86 | 188.9870 | [M – H]<sup>-</sup> | 109.0292 | 99% | -0.51 | Resorcinol monosulfate | Organic acids and derivatives |
| 4.89 | 283.0840 | [M – H]<sup>-</sup> | 107.0498, 175.0253,<br>157.0141, 113.0244,<br>85.0288 | 100% | 0.70 | p-Cresol glucuronide | Organic oxygen compounds |
| 5.18 | 231.0797 | [M – H]<sup>-</sup> | 74.0243, 156.0457,<br>128.0501 | 100% | 0.67 | Indole-3-acetyl-glycine | Organic acids and derivatives |
| 5.20 | 171.1036 | [M – H]<sup>-</sup> | 127.1124, 11.0804 | 100% | -0.78 | 8-oxononanoate | Organic acids and derivatives |
| 5.70 | 297.0994 | [M – H]<sup>-</sup> | 121.0653, 175.0254,<br>113.0242, 85.0287 | 96% | 0.56 | 6-(4-Ethylphenoxy)-3,4,5-trihydroxyoxane-2-carboxylic acid | Organic oxygen compounds |
| **Positive ionization mode** |  |  |  |  |  |  |  |
| 4.66 | 180.065 | [M + H]<sup>+</sup> | 105.0327, 77.0377 | 100% | 0.70 | Hippuric acid | Benzenoids |

DHPPA Dihydroxyphenylpropionic acid; $m/z$ mass to charge ratio; RT retention time; TCA, tricarboxylic acids. \*Corresponds to class assignment. Only those compounds that had MS/MS information have been included.

In plasma, two p-cresol derivatives, two adducts of cholic acid, and twelve glycerophospholipids, including nine phosphatidylcholines and three phosphatidylethanolamines, were identified as significantly different in SHR compared to WKY, while in urine, 14 compounds were identified: one nucleoside, that is pseudouridine; one organoheterocyclic compound, that is, uric acid; four organic acids and derivatives; four organic oxygen compounds; one benzenoid; and two phenylpropanoids and polyketides.

Finally, the results of the pathway enrichment analysis of plasma and urinary metabolites significantly associated with SHR are shown in **Figure 3A** and **3B**, respectively (detailed information is included in the Supplementary Material). These analyses showed significant or borderline significant enrichment of several pathways related to hypertension in plasma, namely steroid hormone biosynthesis (combined *p* value = 0.0049), primary bile acid biosynthesis (combined *p* value = 0.063), and arachidonic acid metabolism (combined *p* value = 0.084) for plasma and purine metabolism in urine (combined *p* value = 0.0071).

**Figure 3.**
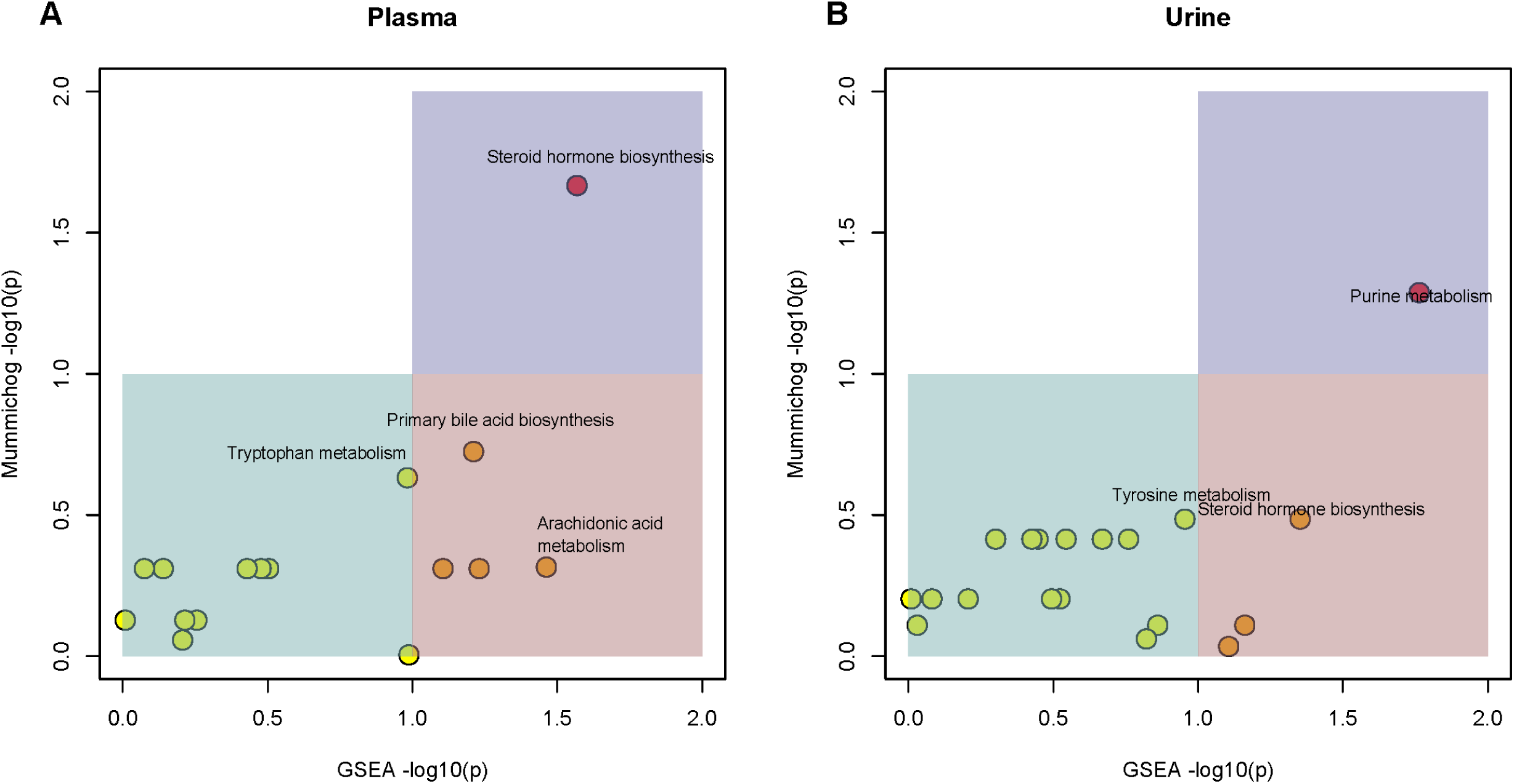
*Mummichog* pathway enrichment analysis of plasma (A) and urine (B) metabolites.

## 4. DISCUSSION

This study deepens the understanding of plasma and urine metabolic differences between 16-week-old hypertensive SHR and aged-matched normotensive WKR rats using LC-qTOF-MS non-targeted metabolomics approaches. Sixteen plasma and 14 urine differential metabolites were semiquantitatively found to be different after deconvolution and were tentatively annotated as a specific compound or as a chemical class. In addition, *mummichog* pathway enrichment analysis has been employed to integrate metabolomic data into biological contexts^18^. The *mummichog* algorithm recognizes specific metabolic pathways associated with altered hypertensive metabolism, mainly steroid hormones from plasma and urine metabolites and purine metabolism from urine metabolites, in consonance with the individual compound annotation. In contrast, gut microbiota alteration and primary liver bile acid metabolism are also associated with hypertension.

The *mummichog* algorithm predicts the functional activities of metabolites and searches for chemical identities by matching the measured mass (*m/z*) of the detected features to a reference metabolic model. This analysis helped us more at the metabolic pathway level without the need to identify individual compounds, and therefore we use it to try to contextualize the changes that we observe and that in the first instance are ‘clouded’ by the presence of other more abundant metabolites like LPCs found in the multivariate analysis. In the case of the sterol hormone biosynthesis pathway, *mummichog* pathway analysis identified several plasma metabolites, indicating that this metabolic pathway is affected in SHR. Specific types of hypertension caused by different forms of inherited mineralocorticoid pathway defects have been described^19,20^. In addition, a previous study reported that serum steroid hormone, progesterone, corticosterone, and cortisol concentrations differed significantly between SHR older than 10 weeks and their age-matched WKY rats, suggesting a role of progesterone in the development of hypertension^21^, independent of other sex hormones^22^. Indeed, this metabolic alteration may be implicated in stress-derived hypertension complications^23^. Our data confirm the role of steroid hormones in the development of hypertension, and because of the high morbidity and mortality associated with hypertension and the high number of undiagnosed people^24^, preventive hypertension follow-up should be implemented in patients with complications in the biosynthesis of steroid hormones to avoid the development of that disease and its complications.

*Mummichog* pathway analysis revealed altered purine metabolism in the SHR, including increased levels of urine uric acid, allantoic acid, and (S)-allantoin. Uric acid is the final oxidation product of purine metabolism and is produced by xanthine oxidase, which oxidizes oxypurines, such as xanthine, to uric acid. In most mammals, such as rats, the hepatic uricase oxidizes uric acid to allantoin, which is excreted in the urine. Epidemiological studies have shown a significant association between hyperuricemia and hypertension^25^. The amount of urate in the blood depends on lifestyle factors such as dietary intake of purines, alcohol, and/or fructose, and physical activity, but it is primarily attributed to genetic factors that determine the level of endogenous urate biosynthesis and the rate of uric acid excretion. Although the underlying molecular mechanisms are not yet well known, it seems that a reduction in endothelial nitric oxide production and stimulation of the vascular renin-angiotensin system, which increases angiotensin II production and subsequently vascular smooth muscle cell proliferation and oxidative stress, may mediate the hypertensive effect of uric acid^26^. The present data confirmed the alteration in purine metabolism during hypertension that may be detected in urine samples, also recommending preventive hypertension-follow-up in people with increased uric acid levels.

In the present study, an elevation of citrate was found in the urine samples but not in the plasma, suggesting that excretion may be a consequence of altered renal metabolism instead of systemic metabolism. Aconitase converts citrate into isocitrate and is inhibited by uric acid; therefore, the elevation of uric acid that we found in SHR might be responsible, or at least contribute, to the accumulation of citric acid in renal cells, which flows out of the mitochondria to the cytosol and is then excreted in urine. In contrast, other authors have reported lower excretion of citric acid in urine from 11-20 weeks old SHR compared with age-matched WKY rats and proposed that it might be related to increased citric acid reabsorption by renal proximal tubules^27–29^. For these reasons, a conclusion regarding citric acid metabolism in SHR cannot be reached, and more investigations are needed to determine its relationship with hypertension.

Higher amounts of urine pseudouridine were found in the SHRs. Pseudouridine is the most widely distributed post-transcriptionally modified nucleotide formed by pseudouridine synthases in RNA. It is involved in the epigenetic regulation of mRNA translation, affects RNA structure and RNA-protein interactions, and thus influences many steps of mRNA metabolism that regulate gene expression^30^, including mitochondrial gene expression^31^. In addition, pseudouridine cannot be degraded in the body; thus, its urinary excretion reflects cellular RNA turnover as well as the glomerular filtration rate. This data agrees with the increased creatinine clearance previously reported in SHR. These authors reported that the glomerular filtration rate progressively increased until 15 weeks old, then reached a plateau and followed by a decline at 60 weeks. Evidence has implicated mitochondria in the most common genetically determined metabolic diseases through different epigenetic mechanisms involved in the modulation of host cell responses to metabolic stress^32^. This is also the case for different mtDNA haplotypes in SHR, which may promote the progression of diverse cardiometabolic phenotypes^33^. Furthermore, the knowledge of the role of mitochondria in the regulation of the smooth muscle phenotype and differentiation is increasing^34^, and pseudouridine has been proposed as a potential biomarker of cardiac dysfunction^31,35,36^. However, to the best of our knowledge, no evidence has connected the pseudouridine epigenetic mechanism with the pathogenesis of hypertension, and future studies are needed in this field.

Accumulating evidence has suggested a key role for the gut microbiota in essential and experimental hypertension in animals and humans^37–40^. A significant decrease in microbial richness, diversity, and evenness, as well as an increased Firmicutes/Bacteroidetes ratio, has also been detected in hypertensive animals and patients^41^. In addition, Vemuri et al.^42^ demonstrated that hypertension contributes to gut microbial translocation and, ultimately, to unhealthy shifts of the gut microbiota. In fact, the protective effects of different antihypertensive peptides have been attributed, at least in part, to their ability to modulate gut microbiota dysbiosis^43,44^. In SHR rats, significant differences in gut microbial composition have been reported compared with WKY rats, specifically regarding the relative abundance of lactic and butyric acid-producing bacteria^45^. Blood pressure regulates in SHR following cross fecal transplantation from WKY, while normotensive animals became hypertensive when receiving fecal transplantation from SHR animals^45^. Our results showed differences between dietary-derived gut microbial metabolites, namely hippuric acid and diverse sulfated and glucuronidated products, in hypertensive and normotensive rats fed the same diet.

Cholic acid is a discriminant metabolite that is lower in the plasma of SHR than in WKY rats. In addition, *mummichog* algorithms have recognized the biosynthesis of primary bile acids as a metabolic pathway affected in SHR, including four taurine-derived salts. Hepatic cholesterol is catabolized into primary bile acids and conjugated to an amino acid side chain to form amphipathic bile salts. Upon transit through the hepatobiliary tract and small intestine, the intestinal microbiota enzymatically transforms bile salts into secondary bile acids. Other authors have previously described the negative role of bile acid metabolism in promoting hypertension in rats^46^. In line with our results, these authors suggested a protective effect of taurine-conjugated bile acids against the development of hypertension and reported that bile acid conjugation was inversely associated with SBP. They described a distinct clustering of taurine-conjugated bile acids that was less abundant in genetically hypertensive Dahl rats than in normotensive animals, independent of salt consumption. Furthermore, the accumulation of microbiota-derived taurine-conjugated bile acids was associated with lower blood pressure, as well as with specific microbiota taxa^46^. Regarding tyrosine metabolism, p-cresol conjugates were lower in hypertensive SHR than in WKY rats, specifically plasma levels of p-cresol sulfate and glucuronide and urine p-cresol glucuronide. P-cresol is mainly generated as an end-product of tyrosine and phenylalanine metabolism by anaerobic intestinal bacteria, followed by secondary conjugation with glucuronide (mainly) and sulfate, to form their corresponding metabolites in the colonic mucosa and liver^27^. In addition, a hypotensive effect of taurine in SHR, associated with an increase in p-cresol derivatives, has been observed, which may be mediated through the modulation of intestinal microbiota metabolism^27^. In addition, we also found reduced levels of urinary hippuric acid, another gut microbiota-derived metabolite^47^, in agreement with data described by Akira et al.^10^ using an NMR-based metabonomic approach. Although one limitation of our study is that we could not analyze the gut microbiota, our data suggest that gut microbiota dysbiosis is associated with hypertension, going beyond, and identifying derived primary liver bile acid metabolism as affected during hypertension.

Although other authors have reported changes in the metabolic profiles associated with hypertension in animal models and humans^35^, the relationship with glycerophospholipids is controversial and depends on the type of lipid^8^. In 2022, Liu et al. identified eight blood pressure-related plasma phospholipids (six phosphatidylethanolamines and two phosphatidylcholine) with predictive value for hypertension risk^48^, while other diacyl-phosphatidylcholines (C38:4 and C38:3) were associated with enhanced hypertension complications^49^. On the other hand, Onuh and Allani^8^ reported that higher levels of different acyl-alkyl-phosphatidylcholines (C42:4 and C44:3) were associated with lower fatal hypertension in humans, possibly due to the protective antioxidant and inflammatory activities of these metabolites. In this work, we identified lower plasma levels of some lysoglycerophospholipids than those in normotensive rats. Decreased levels of lysophospatidilcholines have been observed in several inflammatory-based diseases, including pulmonary arterial hypertension, and were associated with an increased mortality risk^50^. However, Jiang et al.^51^ proposed lysophospatidilcholines as a biomarker of hypertension, suggesting that the process of LDL oxidation may induce their generation, which interferes with nitric oxide production and therefore induces hypertension^51^. Therefore, further investigations in humans with hypertension are needed to confirm certain assumptions related to glycerophospholipid metabolism in those patients.

## 5. CONCLUSION

The investigation of metabolites related to blood pressure regulation is of great interest to clarify the disease mechanisms and identify new biomarkers for early detection and diagnosis before the onset of hypertension, thus avoiding further complications in individuals who are at risk of hypertension. Our research suggests an association of gut microbiota, bile acids, steroid hormones, and purine metabolism with the development of hypertension and its complications. Based on our data, recommending preventive hypertension-follow-up in people with altered steroid hormones metabolism, increased uric acid levels, and dysbiosis may be a good strategy for preventing hypertension in these risk populations.

## Acknowledgments

RICORS funded by the Recovery, Transformation and Resilience Plan 2017-2020, ISCIII, and by the European Union – NextGeneration EU, ref. RD21/0012/0008, and RICORS funded by the Recovery, Transformation and Resilience Plan 2021-2024, ISCIII, and by the European Union – NextGeneration EU, ref. RD24/0013/0007.

## Supporting information

**Supplemental Table 1.** Pathway enrichment analysis of plasma and urinary metabolites significantly associated with SHR.

